# IDRWalker: A Random Walk based Modeling Tool for Disordered Regions in Proteins

**DOI:** 10.1101/2024.03.17.585378

**Authors:** Guanglin Chen, Zhiyong Zhang

## Abstract

**Motivation:** With the advancement of structural biology techniques, the elucidation of increasingly large protein structures has become possible. However, the structural modeling of intrinsically disordered regions in proteins remains challenging. Particularly in the case of large protein complexes, it is difficult to rapidly construct models for all intrinsically disordered regions using existing methods. In the nuclear pore complex, a gigantic protein machine of interest, intrinsically disordered regions play a crucial role in the function of the nuclear pore complex. Therefore, there is a need to develop a modeling tool suitable for intrinsically disordered regions in large protein complexes.

**Results:** We have developed a program named IDRWalker based on self-avoiding random walks, enabling convenient and rapid modeling of intrinsically disordered regions in large protein complexes. Using this program, modeling of all disordered regions within the nuclear pore complex can be completed in a matter of minutes. Furthermore, we have addressed issues related to peptide chain connectivity and knot that may arise during the application of random walks.

**Availability and implementation:** IDRWalker is an open-source Python package. Its source code is publicly accessible on GitHub (https://github.com/zyzhangGroup/IDRWalker).

## Introduction

Large protein complexes play crucial roles in many biological processes, but the structural elucidation of such a complex is a challenging task. Cryo-electron microscopy (cryo-EM) is currently one of the most widely used techniques for resolving the structures of protein complexes^*1, 2*^, which reconstructs 3D electron density maps with atomic-level resolution. However, the high resolution density map is often limited to well folded domains within a complex, with the flexible regions remaining ambiguous^*3*^. Additionally, difficulties in sample preparation and issues such as sample heterogeneity make it challenging to obtain high-resolution structures for protein complexes^*4*^. The larger the structure that needs to deal with, the higher the likelihood of encountering these issues, making it difficult to achieve a high-resolution density map.

The emergence of integrative modeling tools such as the Integrative Modeling Platform (IMP)^*5*^, Haddock^*6*^, and Assembline^*7*^ has provided a new approach for building structural models of a large protein complex using a relatively low-resolution density map. The idea involves first individually solve the high-resolution structure of each protein subunit. Subsequently, guided by density map and information from other experiments^*8*^ such as nuclear magnetic resonance (NMR)^*9*^, cross-linking mass spectrometry (XL-MS)^*10*^, fluorescence resonance energy transfer (FRET)^*11*^, and others, the complete structure of the complex is assembled. This approach has already achieved success in large systems like the nuclear pore complex (NPC)^*12*^ with a molecular weight of ~120 MDa. In recent years, with the assistance of machine learning-based structure prediction methods, such as AlphaFold2^*13*^ and RoseTTAFold^*14*^, high-cost experiments are no longer the exclusive means to obtain accurate structures for each subunit. AlphaFold2 can now yield results comparable to experimental methods, significantly reducing the costs associated with applying integrative modeling to large protein complexes^*15*^.

Despite the aforementioned progresses, the challenge remains when dealing with intrinsically disordered regions (IDRs) in proteins^*16*^. Since it is difficult to capture the dynamic nature of IDRs by conventional experimental methods^*17, 18*^, modeling methods become indispensable for predicting structures in these regions. Although some methods, including multiple sequence alignment (MSA)^*19*^, well-designed score functions^*20-22*^, and neural networks^*13, 14*^, have applied to enhance modeling results of well-folded proteins, they are not efficient for IDRs. Therefore the above tools, though intended to refine results, paradoxically fails to significantly improve predictions in these IDRs^*23*^ and instead increases the computational effort.

Compounding this challenge is a daunting task when dealing with large protein complexes. As a system increases in complexity and size, the burden of additional calculations becomes increasingly unbearable^*24*^. Although treating a large number of atoms as a rigid body is commonly used to reduce computational load when modeling a large complex^*7*^, it is not suitable for IDRs with high flexibility. Considering these issues, in this work, we propose a new approach: ensuring efficiency by modeling with a simple algorithm, which in this paper is self-avoiding random walk. Although the quality of the model thus obtained may not be good enough, further improvement can be achieved in other ways such as molecular dynamics (MD) simulations^*17*^.

Although it is simple, the random work method becomes increasingly burdensome to implement as the complexity of the system increases. Therefore, we have written a Python package called IDRWalker that automates the process of modeling IDRs in proteins by random walk. We have tested this tool by several protein complexes. In the human NPC, we examined the running speed of the IDRWalker when applied to a gigantic protein machine. In the calcium channel protein, we verified the connectivity of the peptide chains in the modeling results. And in the SARS-CoV-2 Omicron spike protein, we investigated how to avoid knots in the models.

In this paper, we will first introduce the implementation of IDRWalker and some technical details. Then, the application of the program in several protein complexes will be presented. Finally, we will discuss the prospect and future improvement of the method.

## Methods

### Overview of IDRWalker

Figure 1 depicts the workflow of IDRWalker. First, the program reads sequence files and structure files, marking the positions of missing regions. Then, a residue generation loop is continuously repeated generating residues one by one until the structures of all missing regions are constructed. Finally, the results are refined and output to the structure file.

**Figure 1.**
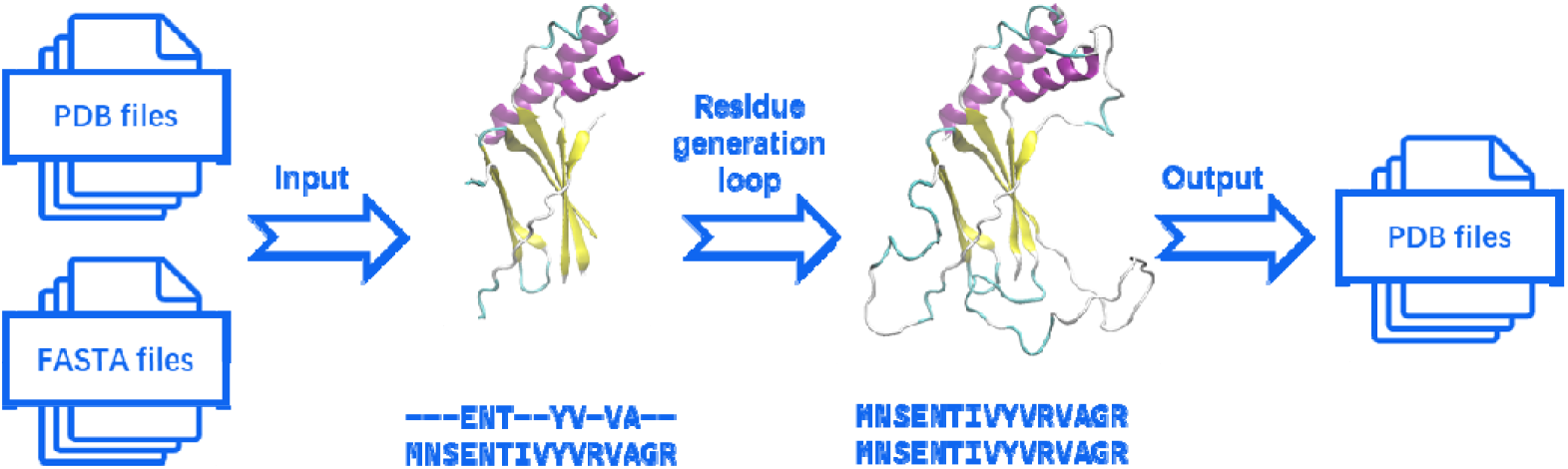
The workflow of IDRWalker

### Input files parsing

The modeling process of random walk is independent across different chains, so when reading input files, it is processed chain by chain. For each chain, the program, based on the read sequence, creates a chain containing the complete sequence by copying the amino acid residue templates which are prepared in advance according to the AMBER ff19SB force field^*25*^. Subsequently, according to the residue numbers, the 3D coordinates in the input PDB file are written to the newly created chain.

To address spatial conflicts in the modeling process at a lower cost, we partition the space where the model exists into grid points. A matrix, referred to as the occupancy matrix in the subsequent text, is used to record whether each grid point is occupied by an atom. Before the modeling begins, the coordinates of each chain are written into the occupancy matrix. It is worth noting that IDRs are typically expansive, meaning that a very large occupancy matrix is required, but a significant portion of the matrix is empty. To avoid the memory wastage issue from the large occupancy matrix, we apply periodic boundary conditions to the occupancy matrix. This allows IDRs to extend beyond the boundaries, enabling the use of a smaller occupancy matrix.

### Residue generation loop

The residue generation loop is the core module of IDRWalker, and its function is to generate the coordinates of a new residue from a residue next to a missing region. The implementation of this module can be divided into the following steps:

#### The coordinates of backbone atoms

Starting from a known residue, the coordinates of a new atom can be generated using the following equations: 

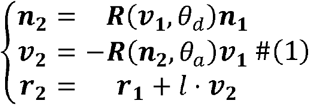

The formula involves a rotation matrix, ***R* (*axisθ*)**, where θ is the angle of rotation chosen around the ***axis***. This matrix can be computed using the Rodrigues rotation formula. The meanings of the other terms are indicated in Figure 2a. In addition to the known residue, the formula also requires bond lengths, angles, and dihedrals. In protein structures, the bond lengths and bond angles are less variable, which can therefore be given directly, for example, using reference values in the AMBER ff19SB force field^*25*^. In contrast, the dihedrals can be rotated relatively free but with some regularities that can be depicted by the Ramachandran plot^*26*^. The dihedral □ in the diagram consists of C-N-C_α_-C, and the dihedral ψ consists of N-C_α_-C-N. Dihedral combinations located within the permissive region of the Ramachandran plot do not lead to spatial conflicts between neighboring residues. There is also a dihedral ω consisting of C_α_-C-N-C_α_ in the main chain, and this dihedral is limited by the peptide bond and takes a limited range of values, usually around 180° and in a few cases around 10°. Based on the described pattern, generating a suitable set of dihedral angles φ, ψ, and ω randomly, and iterating Equation 1 for three times will yield the coordinates of the main chain atoms N, C_α_, and C for the next residue. During the iteration process, it is essential not only to record the generated atomic coordinates but also to keep track of the normal vectors ***n*** and direction vectors ***v*** for use as inputs in the next iteration. Due to issues such as numerical precision, the lengths of the normal vectors and direction vectors may slightly decrease after each iteration. The accumulation of errors over multiple iterations can lead to a modeling failure. For this reason, it is necessary to normalize the normal vectors and direction vectors after a certain number of iterations to mitigate the impact of accumulated errors.

**Figure 2.**
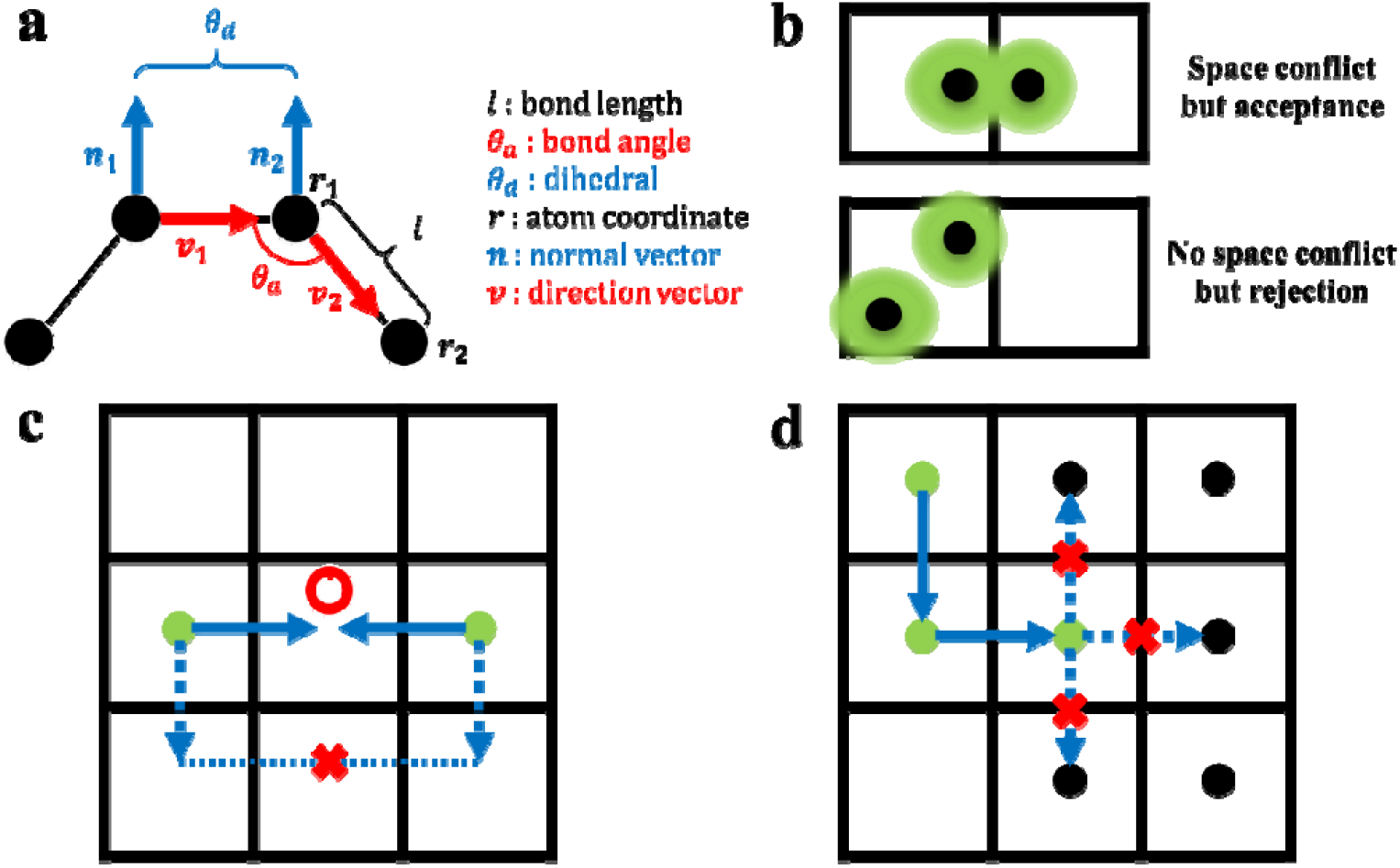
Some details in the residue generation loop. (a) The geometry of four atoms. (b) possible misjudging of space conflict. (c) The connectivity of missing regions. (d) A dead-end of self-avoiding random walk.

#### The coordinates of side chain atoms

Random walk only generates coordinates for the main chain atoms C, C_α_, and N, remaining the coordinates of the side chain atoms unknown. Since the C_α_ atom is at the center of a tetrahedral configuration, the orientation of the side chain can be determined given the coordinates of the main chain atoms as well as the optical activity. The latter is implicit in the amino acid template. Accordingly, the coordinates of the side chain atoms can be obtained as follows: move the amino acid template with Kearsley algorithm^*27*^, aligning its main chain with the newly generated main chain atoms. Subsequently, write the coordinates of the template’s side chain atoms into the new residue.

#### The oxygen atom in a peptide bond

Random walk also does not provide the coordinates of the oxygen atom in a peptide bond. The O atom in the peptide bond forms a double bond and is therefore co-planar with the neighboring C, C_α_, and N atoms, thus its coordinates can be expressed as a linear combination of the coordinates of these three atoms. If using the bond lengths and angles from the AMBER ff19SB force field^*25*^, the coordinates of O atom can be calculated using the following formula: 

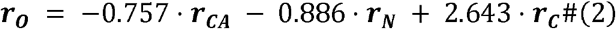

Due to the involvement of different atoms in bonding, the coordinates of the C-terminal oxygen atom cannot be calculated using the Equation 2. It can be assumed to lie in the same plane as the N, C_α_, and C atoms of the same residue, and then computed using the following formula: 

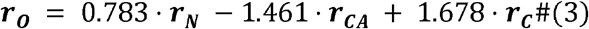

Unlike the situation where atoms determining the side-chain position are within the same residue, the atoms related to the position of the oxygen atom in the peptide bond are in different residues. Due to this issue, the coordinates of the oxygen atom cannot be immediately determined after generating a new residue, they can only be finalized after modeling the entire chain.

#### Spatial conflicts check

Once the coordinates of all heavy atoms in the new residue, except for the oxygen atom in the peptide bond, are determined, it is crucial to check their reasonability. The first step is to check for spatial conflicts. By comparing coordinates with the occupancy matrix and observing whether any atoms are positioned within already occupied grids, it is possible to make a rough assessment of spatial conflicts. Compared to calculating pair distances between atoms, this method involves significantly less computational effort. However, it can only eliminate cases of large-scale overlap and may not entirely rule out the possibility of spatial conflicts. Additionally, there is a risk of misjudging reasonable configurations as spatial conflicts (Figure 2b). If there is a misjudgment involving the main chain atoms of adjacent residues, it could introduce false constraints to the main chain dihedrals. Considering that spatial conflicts between adjacent residues have already been addressed through the Ramachandran plot, when evaluating spatial conflicts, backbone atoms except C_α_ are excluded.

#### Connectivity check

To model a missing region in the middle of the peptide chain with random walk, it’s essential to make sure that the two ends of the missing region intersect at the same place after a certain number of random walk steps (Figure 2c). We used Monte Carlo method to study the random walk law of protein main chain. By counting the walk of 100 residues for 1000 times, we found that when the number of steps is large, the distribution of the distance from the starting point can be approximated by a normal distribution (Figure S2), and the relationship between and the number of steps can be obtained by linear fitting (Figure S1). When the number of steps is small, although it cannot be described by the normal distribution, it can be directly described by the result of multiple walks. After a residue is generated by a random walk, the probability of connecting the peptide chain can be evaluated according to the above law, only results with a high likelihood of connecting pass the connectivity check.

#### Forward or backward

If a newly generated residue passes the checks above, its coordinates can be written into the occupancy matrix, and the random walk moves forward. Attempts to generate a new residue are not always successful. If a generated residue fails the checks, the process needs to be repeated until success. If there are too many failed attempts, for example, exceeding 1000, which indicates that the random walk might be stuck in a dead-end (Figure 2d). In such a case, moving backward is necessary, and the coordinates of the current residue should be removed from the occupancy matrix.

The above represents all the steps for modeling a single residue. By iteratively running the residue generation loop, modeling for all missing IDRs can be completed. To prevent the modeling sequence from influencing the results, the missing region for generating residues is randomly selected each time during the modeling process.

#### Refinement

Due to deficiencies in the spatial conflict check, a model obtained through the above process needs refinement, which is achieved by performing an energy minimization using the double-precision version of GROMACS^*28*^. The model passed the energy minimization can be utilized for MD simulations.

## Results and Discussion

### The human NPC

The NPC is a crucial structure governing molecular traffic between the nucleus and the cytoplasm. Notably, it is characterized by a significant presence of disordered regions like FG-repeats in its constituent proteins^*12*^. The FG repeats within the central transport channel are primarily contributed by Nup98, Nup54, Nup58, Nup62, etc. These proteins are believed to play a crucial role in controlling the entry and exit of substances into the cell nucleus^*15*^. However, the specific mechanism can be only explained by some hypotheses that have not been completely verified. In order to advance the study of the mechanism, it is essential to establish models for FG-repeats and other disordered regions within the NPC.

The NPC, due to its enormous size, poses significant challenges for modeling. Although IDRWalker automates the modeling process, organizing the input files remains a non-trivial task. We downloaded the structure and sequence files of the human NPC (PDB ID: 7r5k)^*15*^, which consists of a total of 808 chains. Subsequently, we saved the structure and sequence of each chain into separate PDB and FASTA files. These files were then sequentially read by IDRWalker to complete the modeling process. The preprocessing of input files is not integrated into IDRWalker, due to the fact that for different systems it may be necessary to write different scripts to handle it.

Figure 3 shows a model of NPC with all missing regions in the resolved structure. The refined model was coarse-grained using martinize.py. The maximum force experienced by atoms in the system reached up to 1.50226 × 10^14^ kJ · mol^−1^ · nm^−1^, indicating spatial conflicts. After 250 steps of energy minimization in a vacuum, the maximum force decreased to below 1000 kJ · mol^−1^ · nm^−1^ . The result can be utilized as the initial conformation for MD simulations.

**Figure 3.**
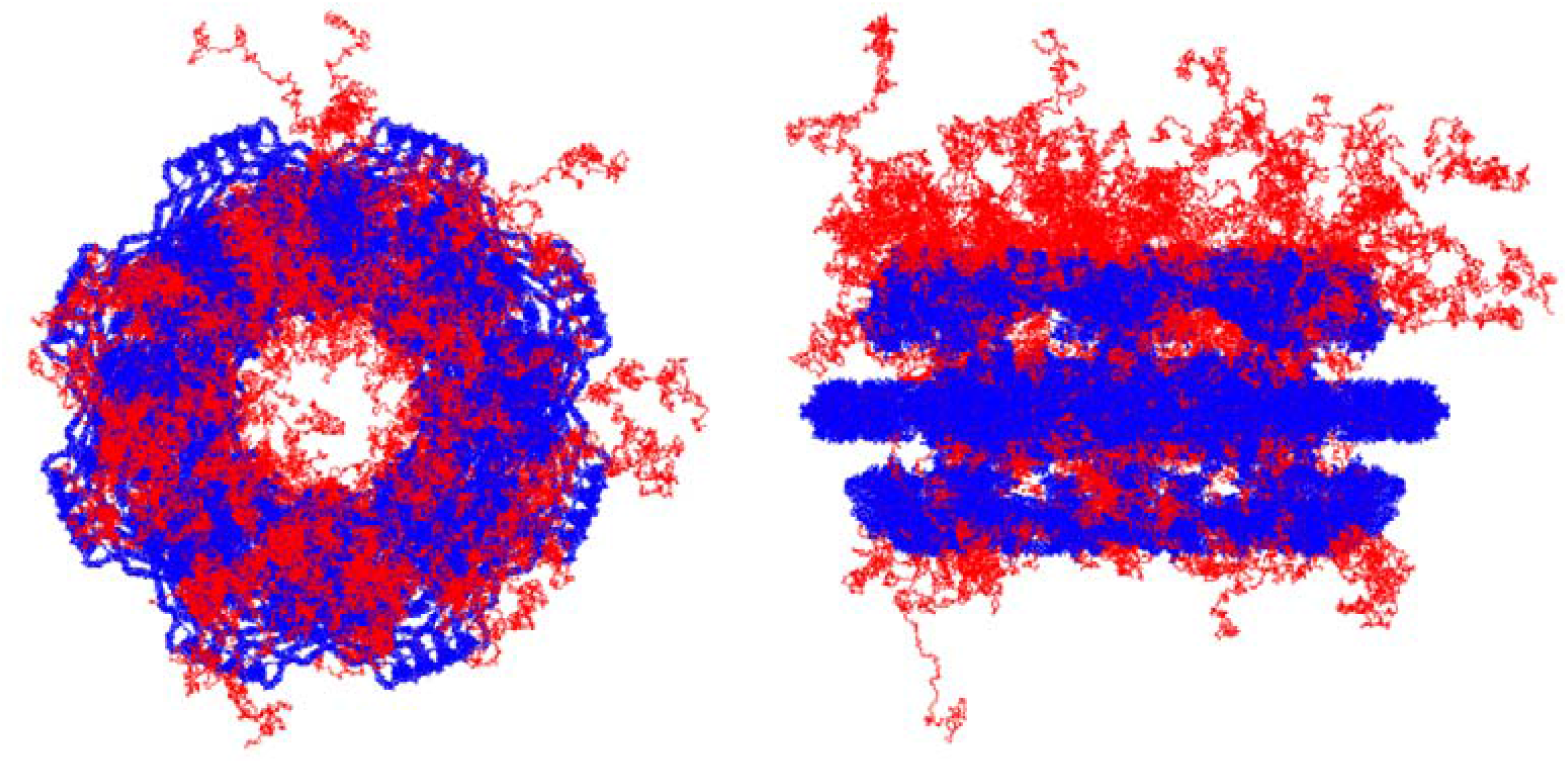
The modeling result of the NPC. Known regions are in blue, modeling regions generated by IDRWalker are in red.

### Calcium channel protein RyR1

The Ryanodine Receptor 1 (RyR1) stands as a pivotal calcium channel protein located predominantly in the sarcoplasmic reticulum of skeletal muscle cells, playing a central role in excitation-contraction coupling^*29*^. Despite its physiological significance, obtaining a comprehensive and high-resolution structure of RyR1 has posed a significant challenge. While the resolved structures have provided valuable insights, certain regions are notably absent in each of the four symmetrical parts of RyR1, with some deletion regions exceeding 100 residues in length. In the structure of RyR1 with PDB ID 8seu^*30*^, there are four gaps in each chain, and the number of missing amino acid residues in each gap are 115, 285, 38 and 22. These gaps in the structural data often encompass disordered or highly flexible segments, however, whether IDRWalker can be used to model the missing regions remains a question because the algorithm in IDRWalker, which ensures peptide chain connectivity, only considers cases within 100 residues.

To verify the effectiveness with gap lengths exceeding 100 residues, we utilized a symmetrical chain from the RyR1 structure as a template for modeling the internal gaps with IDRWalker. The modeling result (Figure 4a) demonstrates that the algorithm remains effective even when the missing region exceeds 100 residues.

**Figure 4.**
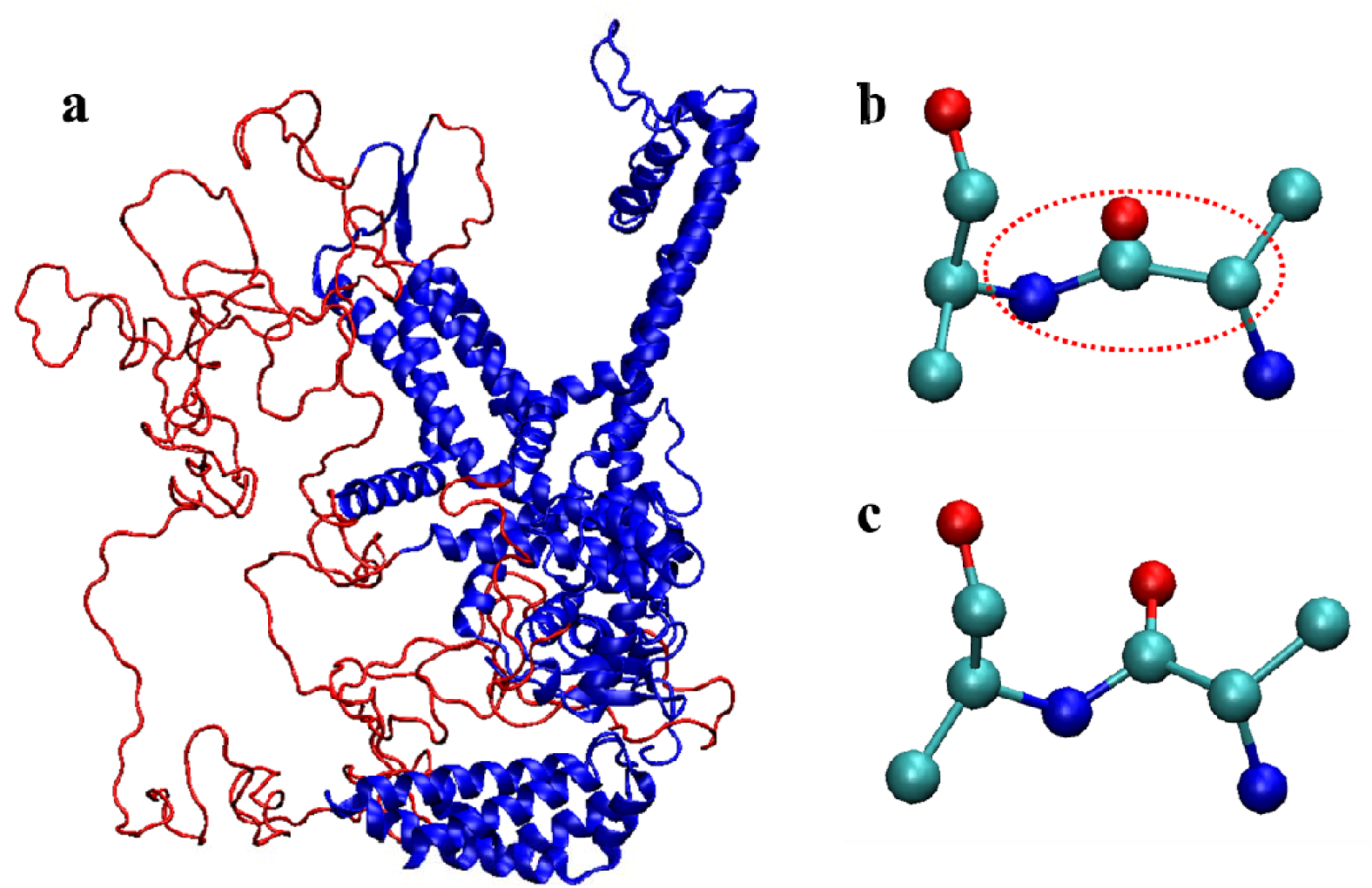
The modeling of A chain in RyR1. (a) The modeling result, the known regions in blue, and the modeling regions in red. (b) Incorrect bond angles and oxygen atom coordinates. (c) The corrected angles and oxygen atom coordinates after energy minimization.

In the modeling result, issues with bond angles may arise, as a result of this, coordinates of some oxygen atoms could also be wrong (Figure 4b). This is because the algorithms ensuring connectivity impose restrictions only on distances without considering angles. Although the patterns of angle variation with the number of walking steps are similar to the distances, constraining angles significantly reduces the success rate of modeling. Therefore, the problem of bond angle anomalies is not addressed during modeling but is corrected through energy minimization afterward (Figure 4c).

### SARS-CoV-2 Omicron spike protein

The SARS-CoV-2 spike protein serves as a key player in the viral infection process, playing a central role in facilitating the virus’s entry into host cells. Comprised of two subunits, S1 and S2, the spike protein undergoes conformational changes that are crucial for mediating the fusion between the viral and host cell membranes. The receptor-binding domain (RBD) within the S1 subunit directly interacts with the host cell receptor, angiotensin converting enzyme 2 (ACE2), initiating the process of viral entry. Mutations, such as delta and omicron, impact the virus’s affinity for ACE2, alter its transmissibility, and potentially confer immune escape. In the resolved structure of the spike protein (PDB ID:7WK4)^*31*^, many segments are missing, and some of these are near active structural domains, drug targets, and mutation sites. Thus, modeling the missing regions in the spike protein is necessary^*32*^.

When modeling the missing portions based on the resolved structure of the SARS-CoV-2 Omicron spike protein, the orientation of terminal residues and limited space may lead to the occurrence of knots between adjacent chains in the trimeric structure. Due to the rarity of naturally occurring knotted proteins^*33*^, knotting during modeling is generally considered irrational. IDRWalker cannot avoid knots either, but due to the small amount of computation, randomness, and the convenience of adjusting the parameters, it is possible to try a large amount of modeling and select a reasonable result (Figure 5). It’s important to note that in the PDB file, the residue numbers are mapped to the wild-type sequence rather than the mutant sequence.

**Figure 5.**
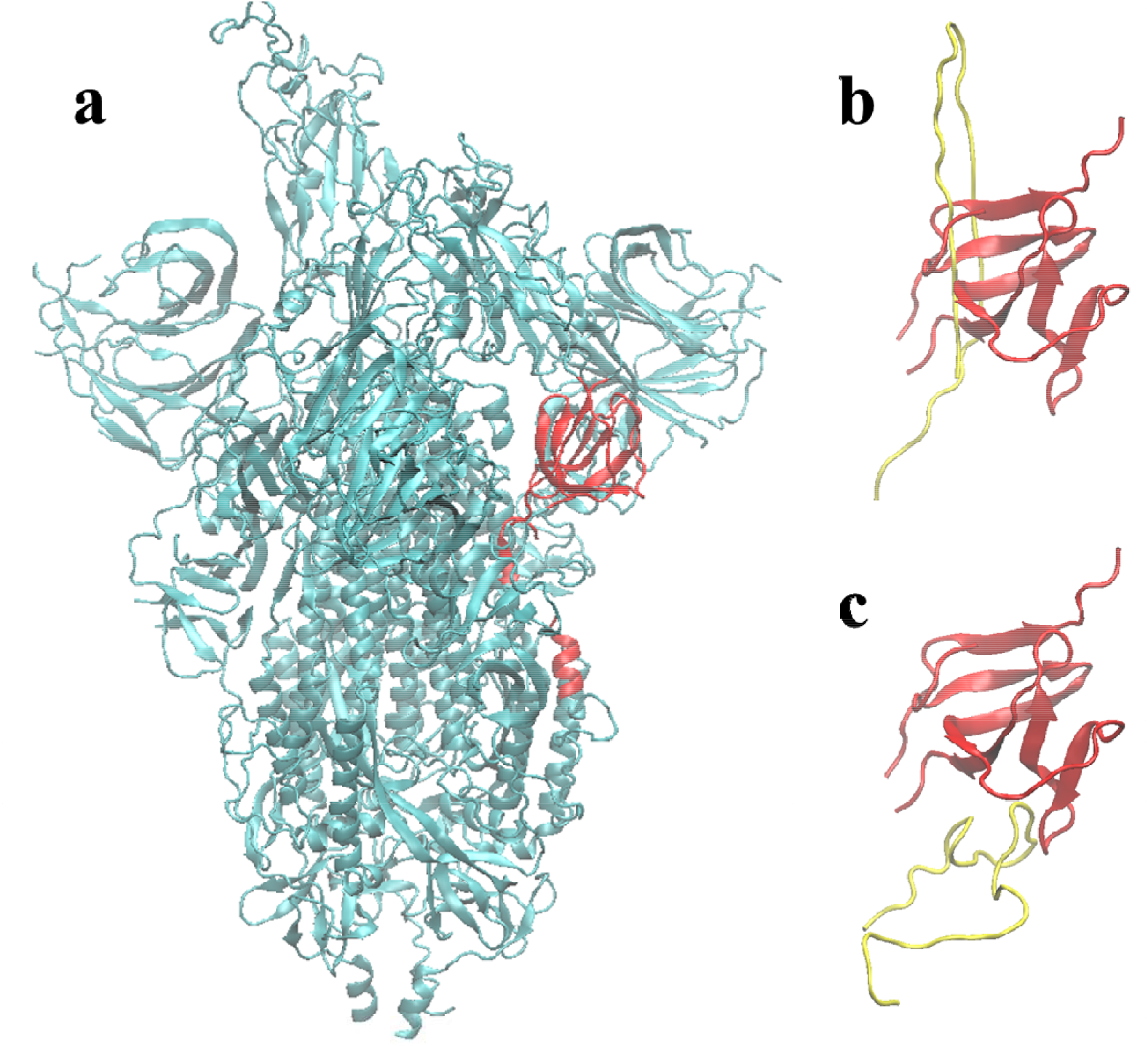
The modeling of Omicron spike protein. (a) The known structure of the spike protein, an area of possible knots is marked in red. (b) A model with knot. (c) A model without knot.

Therefore, it is necessary to map the residue numbers to the sequence of Omicron before modeling.

### Efficiency of the program

The motivation for designing the program is to address the efficiency issue associated with modeling IDRs in complex systems, so it is important for the program to run with sufficiently fast speed. Table 1 displays the time consumption and various parameters for modeling different systems. As observed, IDRWalker can complete the modeling of IDRs with fewer than 500 residues in less than 1 second. Even for a huge protein complex like the NPC, the runtime is only a few minutes. The above results were obtained using only a single CPU core. If parallel processing is employed, it allows the simultaneous generation of multiple different models, thereby further enhancing modeling efficiency. However, when modeling large systems, parallel processing may face significant limitations due to substantial memory usage. This is an area that requires improvement.

**Table 1.** The running details of IDRWalker.

| Modeling regions | PDB ID | Modeling residues | Running time | Memory usage |
| --- | --- | --- | --- | --- |
| Missing regions in human NPC | 7R5K | 248928 | 81 s | 2.21 GB |
| Internal gaps in A chain of RyR1 | 8SEU | 460 | < 1 s | 119.2 MB |
| Internal gaps in 2 neighboring chains of Omicron S protein | 7WK4 | 157 | < 1 s | 119.2 MB |

## Conclusion

Currently, there are challenges in modeling the IDRs of large proteins, including issues related to high computational demands and complex procedures. We have addressed these challenges by employing a random walk method, which allows for modeling the IDRs with low computational costs. Additionally, we have developed a program named IDRWalker to simplify the modeling steps, making it more efficient for application to large biomolecular complexes.

With IDRWalker, we first modeled the IDRs in the NPC. By examining the runtime and modeling results, we have preliminarily validated the feasibility of the program. Then we modeled the IDRs of RyR1 and the Omicron spike protein. According to the results, we verified that IDRWalker is capable of addressing issues that may arise during modeling, such as discontinuities and knots in the peptide chains.

To reduce computational load, the current program does not utilize experimental data. The potential for integrative modeling within the framework of this program is worth exploring. For instance, the density map of disordered regions can be used to control the modeling. Another example is applying the algorithm used in IDRWalker, which ensures chain continuity, to constrain the distance between residues which can be determined through experiments such as NMR, FRET, and XL-MS. We will explore the implementation of these ideas in future research endeavors.

## Acknowledgments

This work is supported by the National Key Research and Development Program of China (2021YFA1301504), the National Natural Science Foundation of China (91953101), and the Chinese Academy of Sciences Strategic Priority Research Program (XDB37040202). We are grateful to Mr. Yundong Zhang for technical support.

## Notes

### Competing Interest Statement

The authors have declared no competing interest.

