## Supplemental Figure 1, Supplemental Figure 2 for "IDRWalker: A Random Walk based Modeling Tool for Disordered Regions in Proteins"

**The PDF file includes:**

Figures S1 to S2

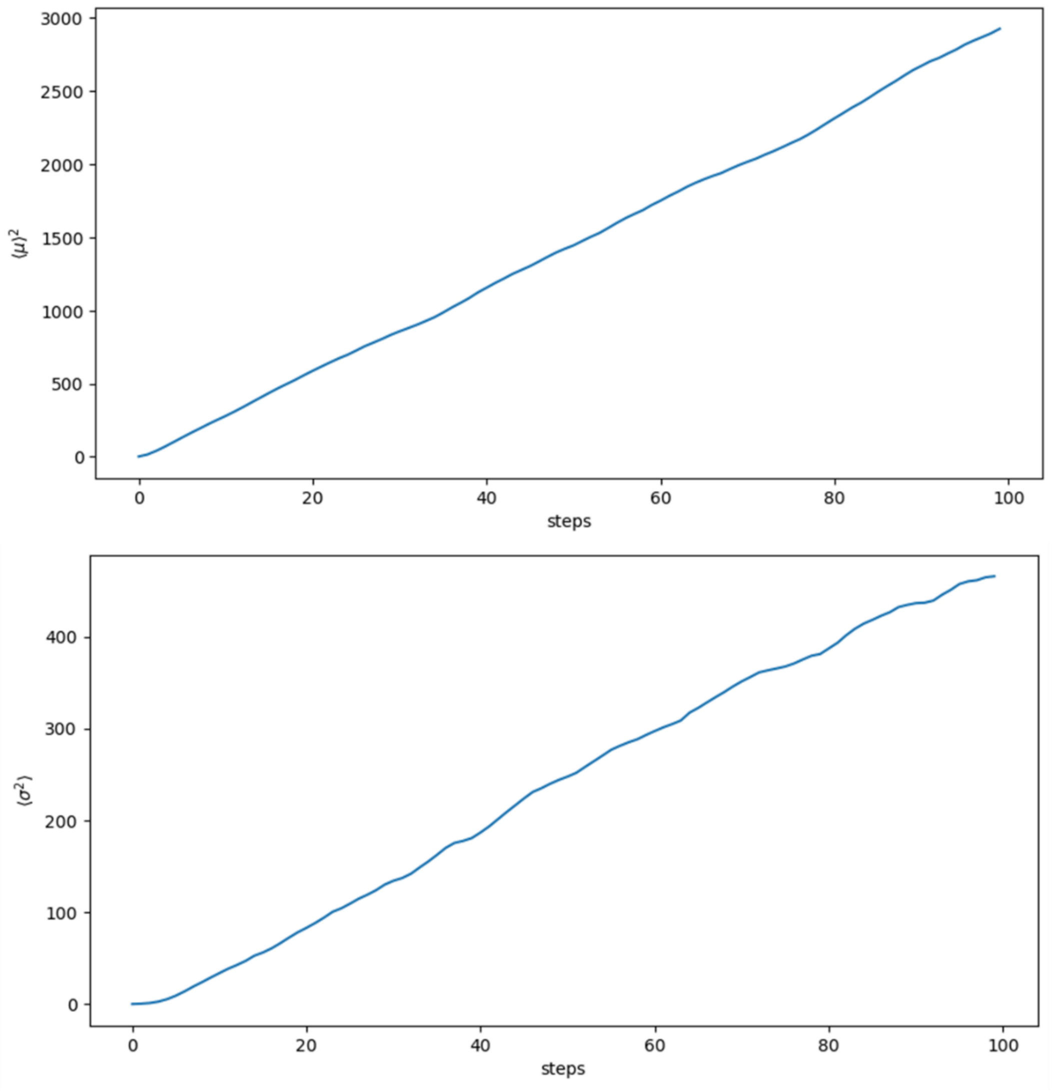


**Figure S1 $\langle\mu\rangle^2$ and $\langle\sigma^2\rangle$ vary with the number of steps**

$\langle\mu\rangle^2$ is the square of the average of the distance from start position during the random walk of the protein backbone. $\langle\sigma^2\rangle$ is the variance of the distance. Statistics from 1000 independent protein backbone random walk paths of 100 residues.

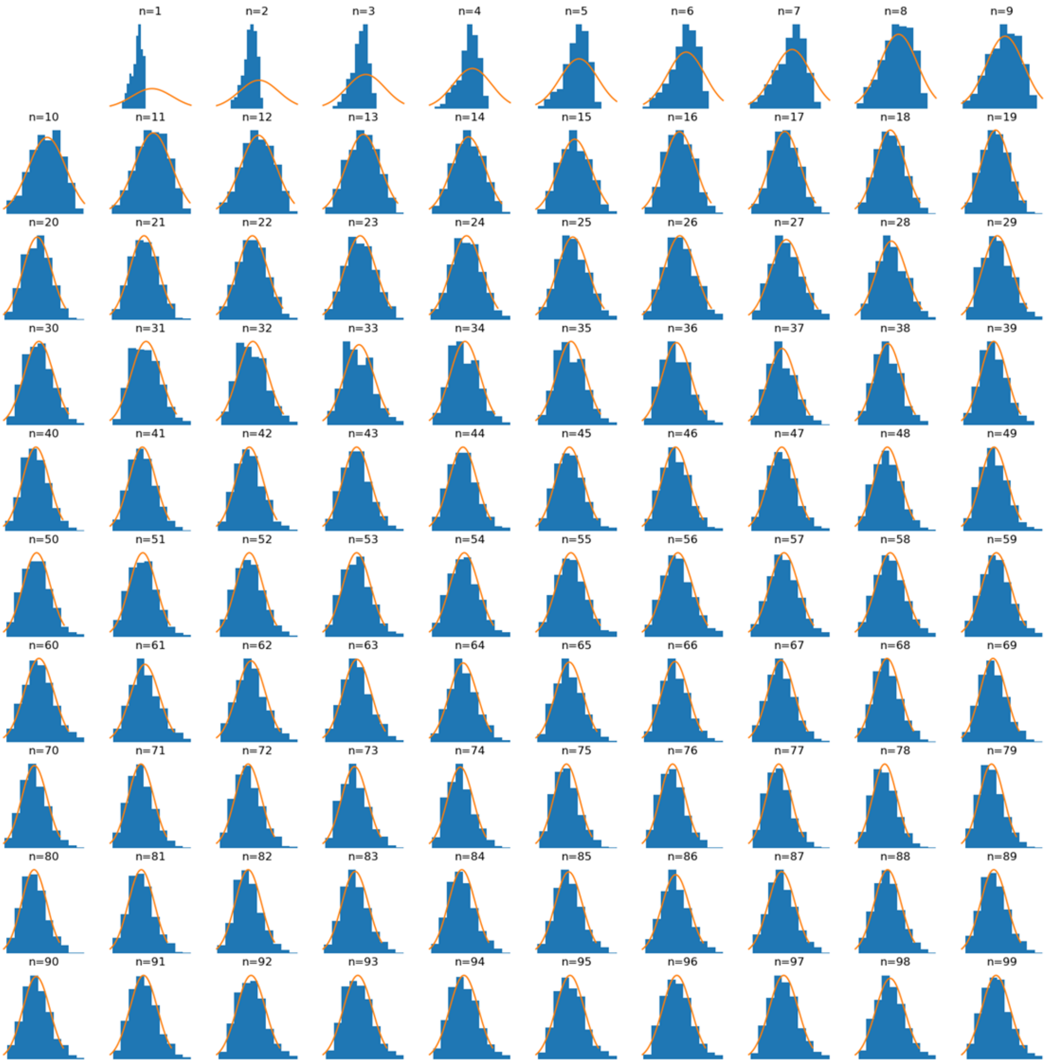


**Figure S2 Approximate description of distance distribution using normal distribution**

Each subplot records the distribution of distances at a specific number of steps. The blue bars are histograms of the distribution of distances, and the orange line indicates the normal distribution used to approximate the distribution of distances, whose $\mu$ and $\sigma$ are obtained from a linear fit. Statistics from 1000 independent protein backbone random walk paths of 100 residues.
